# A diploid chromosome-level genome of *Eucalyptus regnans*: unveiling haplotype variance in structure and genes within one of the world’s tallest trees

**DOI:** 10.1101/2024.06.29.600429

**Authors:** Scott Ferguson, Yoav D Bar-Ness, Justin Borevitz, Ashley Jones

## Abstract

**Background:** *Eucalyptus regnans* (Mountain Ash) is an Australian native giant tree species which form forests that are among the highest known carbon-dense biomasses in the world. To enhance genomic studies in this ecologically important species, we assembled a high-quality, mostly telomere-to-telomere complete, chromosome-level, haplotype-resolved reference genome. We sampled a single tree, the Centurion, which is currently a contender for the world’s tallest flowering plant.

**Results:** Using long-read sequencing data (PacBio HiFi, Oxford Nanopore ultra-long reads) and chromosome conformation capture data (Hi-C), we assembled the most contiguous and complete *Eucalyptus* reference genome to date. For each haplotype, we observed contig N50s exceeding 36 Mbp, scaffold N50s exceeding 43 Mbp, and genome BUSCO completeness exceeding 99%. The assembled genome revealed extensive structural variations between the two haplotypes, consisting mostly of insertions, deletions, duplications and translocations. Analysis of gene content revealed haplotype-specific genes, which were enriched in functional categories related to transcription, energy production and conservation. Additionally, many genes reside within structurally rearranged regions, particularly duplications, suggesting that haplotype-specific variation may contribute to environmental adaptation in the species.

**Conclusions:** Our study provides a foundation for future research into *E. regnans* environmental adaptation, and the high-quality genome will be a powerful resource for conservation of carbon-dense giant tree forests.

## Background

*Eucalyptus* forests are widespread across Australia and extend north into tropical islands. They provide habitat to a rich biodiversity of marsupials, birds and insects, being key foundation species in natural ecosystems [1]. *Eucalyptus* trees are highly diverse and adaptable, exhibiting resistance to extreme droughts, fires and floods. The *Eucalyptus* genus contains over 900 species that have variable genome sizes of approximately 400-700 Mbp [2], high heterozygosity [3] and high frequency of structural variants [4]. With different phenotypes and adaptive traits to varying environments, there is an increasing need for representative genomes.

*Eucalyptus regnans* (Mountain Ash, also known as Swamp Gum and Stringy Gum) is part of the diverged subgenera *Eucalyptus*, formerly known as *Monocalyptus* with several sections and 100 species, including the alpine specialist snow gum, *E. pauciflora,* that is facing dieback [5]. Representing a grove of giant trees in Tasmania, *E. regnans* forests are among the highest known carbon-dense biomasses in the world [6]. They annually sequester and store large amounts of carbon, with the wet temperate forests having the highest above and below ground carbon densities of 1,000 tC/h [7] to 1,312 tC/h [6]. This highlights their significance in mitigating climate change, but they are also under threat from climate change, particularly widespread bushfires that can occur in Australia [8, 9]. Increased logging and deforestation is also becoming a widespread concern. Therefore, there is an increasing need for conservation, management, and restoration of these forests.

Among these forests, the *E. regnans* tree known as “Centurion” (Figure 1), is currently a pre-eminent candidate for the world’s tallest known flowering plant [10]. Captivating researchers and enthusiasts alike, it has been measured at 99.6 m by an aerial laser scanning LIDAR forest inventory [11] and at 99.8 m by tape drop techniques [12]. In 2018, the tree was remeasured at 100.5 m using ground-based observations with a Laser Technologies TruPulse 360 [13]. Still alive and growing, despite being partially burnt in a bushfire, it is among the tallest known angiosperms [14, 15]. Currently, the tallest known trees are the non-flowering *Sequoia sempervirens* (coast redwoods, such as the “Hyperion”), which can achieve over 112 m in height [16]. Further research into the world’s tallest trees provides valuable opportunities to understand tree growth, carbon sequestration and wood production.

**Figure 1.**
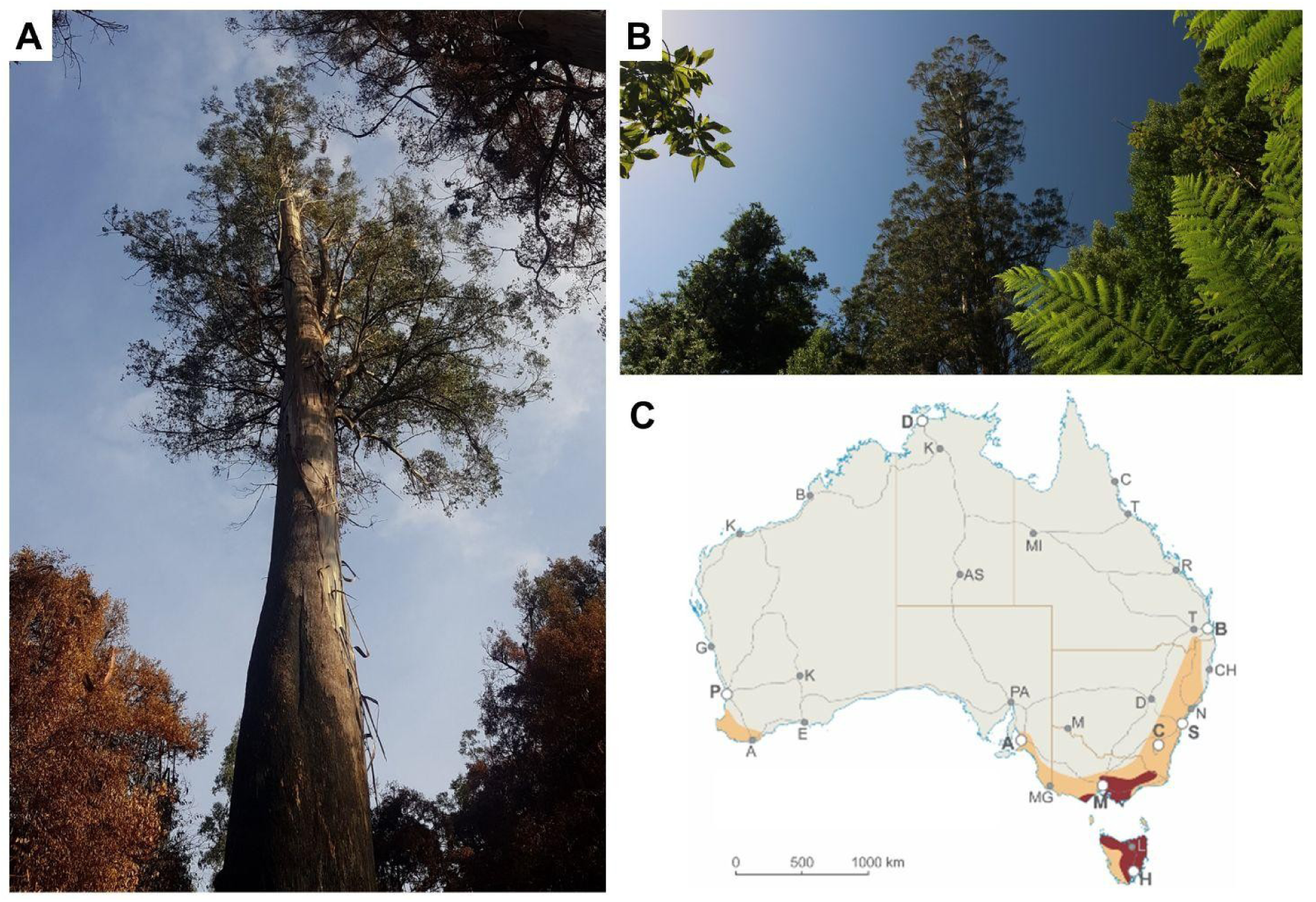
*Eucalyptus regnans*, the Centurion, located in Tasmania, Australia. (A) Picture is of the Centurion after national bushfires in Australia. (B) The Centurion before the bushfires. Photographs by Yoav Daniel Bar-Ness (Giant Tree Expeditions). (C) Species occurrence distribution map of *E.regnans* in Australia. Red represents natural distribution, orange represents low frequency plantations. Alphabetical letters represent major cities in Australia, bold is capital cities. Key areas of natural distribution are Hobart (H), Launceston (L), and Melbourne (M). Map sourced from [21], being provided by the author Dean Nicolle.

To enable further studies into *E. regnans* and carbon-dense giant trees, we assembled a haplotype-phased chromosome-level genome of the Centurion, using Pacific Biosciences (PacBio) HiFi reads, Oxford Nanopore Technologies (ONT) ultra-long reads and Hi-C chromosome conformation capture, in the hybrid assembler Hifiasm ultra-long (UL) [17]. Long-read sequencing technologies now enable complete, telomere-to-telomere (T2T) assemblies of complex genomes, such as the human genome [18], kiwifruit [19] and maize genome [20]. Using this approach, we assemble the most complete *Eucalyptus* genome to date, being the first chromosome-level, haplotype phased, diploid assembly, approaching complete T2T quality. Our genome provides insights into the structural variation between haplotypes and enables further studies into genome evolution in *Eucalyptus*.

## Results

### Long-read native DNA sequencing

To assemble the genome of *E. regnans* the Centurion, we extracted high-molecular weight DNA from leaves for long-read sequencing with PacBio for HiFi reads and ONT for ultra-long reads ≥ 40 kb. A portion of leaf tissue was crosslinked for Hi-C chromosome conformation capture followed by short-read sequencing with Illumina. Sequencing generated of 92.07 Gbp HiFi (N50 15.80 Kbp, ∼176x coverage), 23.53 Gbp ONT (N50 46.73 Kbp, ∼49x coverage), and 8.03 Gbp Hi-C (150 bp paired end) (Figure 2.A, 2.B, 2.C). ONT reads were filtered, removing all reads < 20 Kbp and < Q7, leaving 21.34 Gbp (N50 28.35 Gbp, ∼41x coverage). Hi-C sequences were contained in 26.77 million read pairs.

**Figure 2.**
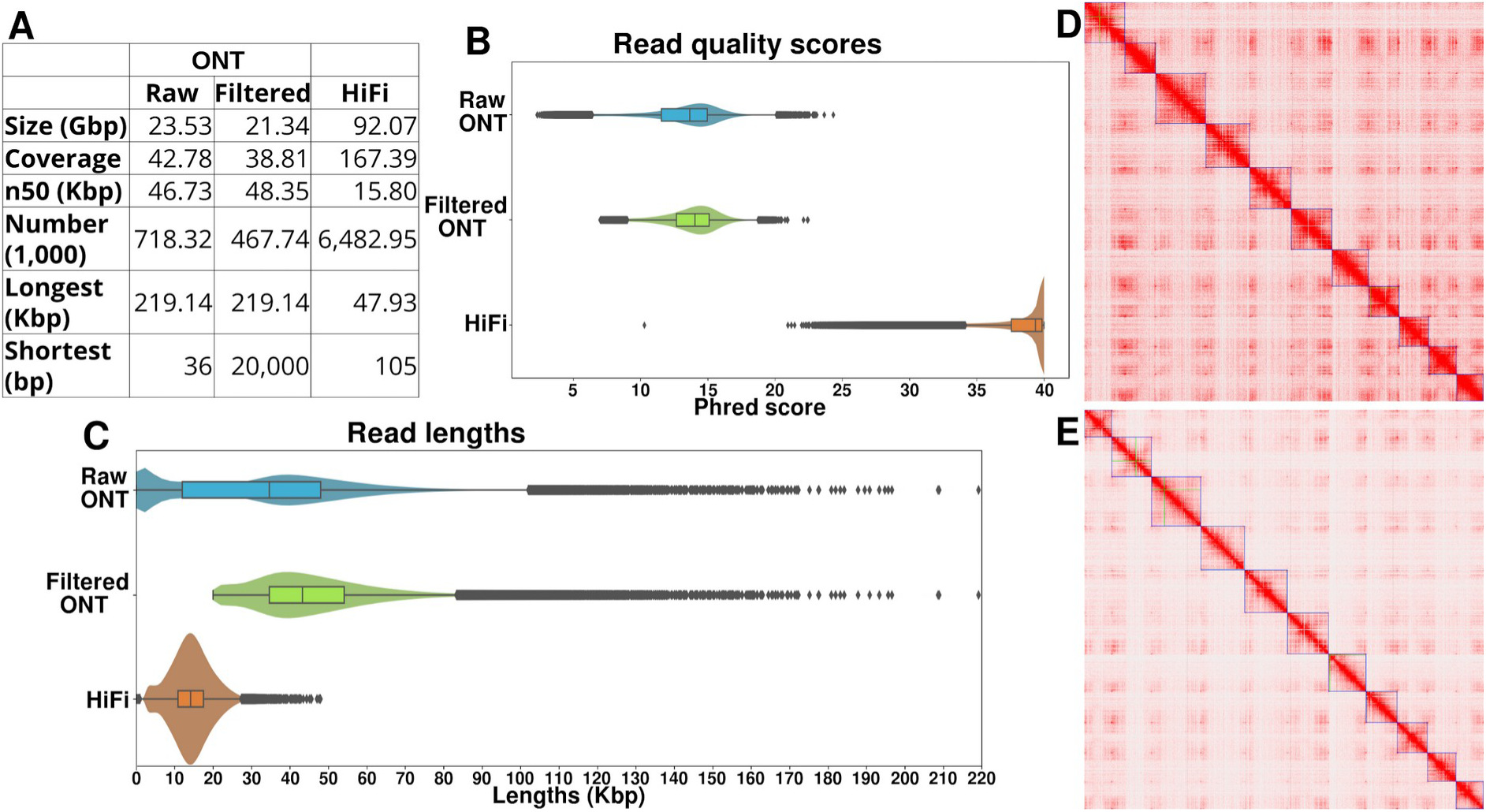
Long-read sequencing statistics and Hi-C contact map. A) Summary statistics for all assembly sequencing data (HiFi, raw ONT, and filtered ONT). B) Violin plot of read quality scores. C) Violin plot of sequencing read lengths. D) Hi-C contact map of scaffolded contigs in *E. regnans* haplotype 1. E) Hi-C contact map of scaffolded contigs in *E. regnans* haplotype 2. Hi-C contact heatmaps were visualised with Juicebox [22].

### Assembly, scaffolding, and telomere identification

Using all data types as input into HiFiasm (UL) (ONT, HiFi, and Hi-C reads), the *E. regnans* genome was assembled into two haplotypes, comprising of 795 contigs 523 Mbp haplotype 1, and 269 contigs 505 Mbp haplotype 2 (Table 1). The Contig N50s for haplotypes 1 and 2 were 36.83 Mbp and 37.75 Mbp, respectively, being the most contiguous contigs for a *Eucalyptus* species to date. After assembly, we investigated our contigs for contamination sequences, finding and removing 18.1 Mbp in haplotype 1 and 17.6 Mbp in haplotype 2. Hi-C reads were independently aligned to both *E. regnans* haplotypes, followed by removal of low-quality aligned reads (MAPQ < 30), chimeric reads, and PCR duplicates. This revealed approximately 2.42 million (9.05%) read pairs contained inter-chromosomal linkage information, and approximately 3.40 million (12.69%) contained intra-chromosomal linkage information (Table 2). Using the linkage information from Hi-C read pairs, both haplotypes were scaffolded into 11 pseudo-chromosomes, representing the correct number of chromosomes (Figure 2.D, 2.E).

**Table 1.** *E. regnans* genome assembly statistics.

|  | **Haplotype 1** | **Haplotype 2** |
| --- | --- | --- |
| **Scaffolded genome Size (bp)** | 523,250,160 | 504,553,124 |
| **Identified telomeres** | 21/22 | 21/22 |
| **% of genome in scaffolds** | 91.60% | 96.07% |
| **Scaffolded N50 (bp)** | 43,985,167 | 48,117,852 |
| **Scaffold count** | 11 | 11 |
| **Contig N50 (bp)** | 36,825,038 | 37,748,426 |
| **Contig L50** | 7 | 7 |
| **Number of contigs** | 795 | 269 |
| **Genome BUSCO complete and (Duplicated)** | 99.27% (2.24%) | 99.14% (1.98%) |
| **Repetitive % (TE %)** | 39.27% (38.16%) | 41.77% (40.67%) |
| **Predicted gene candidates** | 71,726 | 64,961 |
| **Proportion of genome in predicted genes** | 14.09% | 13.67% |
| **Gene BUSCO complete and (Duplicated)** | 97.25% (13.93%) | 95.23% (12.51%) |

**Table 2.** Hi-C chromosome conformation capture linkage statistics.

|  | Haplotype 1 | Haplotype 2 |
| --- | --- | --- |
| Hi-C Read Pairs | 26,768,465 |  |
| Inter-chromosomal | 2,387,828 (8.92%) | 2,456,369 (9.18%) |
| Intra-chromosomal | 3,408,313 (12.73%) | 3,384,634 (12.64%) |
| Uninformative Read Pairs | 20,972,324 (78.35%) | 20,927,462 (78.18%) |

BUSCO analysis indicated high completeness for both haplotypes (Table 1 and Supplementary Table S1). Scaffolding was assessed by aligning both haplotypes to *E. grandis* [23] and against each other, Supplementary Figures S1, S2, and S3. Scaffolding was confirmed and all scaffolds were named according to the *E. grandis* chromosome names, which is the custom for *Eucalyptus*. For both haplotypes, 21 out of 22 telomeres were identified. In haplotype 1, Chromosome 4 was missing the 5′ telomere, while haplotype 2 was missing a telomere from the 5′ end of Chromosome 11. The identified telomere sequence was AAACCCT. This telomere sequence has also been observed in the majority (282 of 332) of Dicotyledons (Magnoliopsida) listed in the telomeric repeat database (https://github.com/tolkit/a-telomeric-repeat-database).

### Genome annotation and gene orthogrouping

Both haplotypes were *de novo* annotated for transposable elements (TE), simple repeats, and genes (Table 1). Repeat annotation resulted in the identification of ∼40.52% of both haplotypes as repetitive, of which ∼39.42% was TE. After soft masking, both haplotypes were annotated for genes. Haplotype-specific HMM models were trained on all available NCBI [24] gene transcripts for *A. thaliana* (Taxonomy ID: 3702) and *Myrtaceae* (Taxonomy ID: 3931). Subsequently, 71,726 and 64,961 genes were predicted for haplotype 1 and 2 respectively.

To examine how similar (or dissimilar) the gene content of the two *E. regnans* haplotypes are, all primary transcripts (longest) were orthogrouped. Orthogrouping places highly similar genes into groups, within and between haplotypes. Genes within each orthogroup are identical genes, gene duplicates or members of the same gene family, and have identical or highly similar function. Of a predicted 125,904 primary transcripts, 97,314 (77.3%) were placed into an orthgroup, the remaining 28,590 (22.7%) were found to be too dissimilar to all other transcripts and not placed within an orthogroup. A total of 86,149 primary transcripts were found to be shared between both haplotypes (haplotype 1: 41,598 haplotype 2: 44,551). The remaining transcripts (haplotype 1: 24,492 haplotype 2: 15,263) were unique to each haplotype (Table 3). Orthogrouping created 39,959 groups, of which 36,882 (95.4%) were shared.

**Table 3.** *E. regnans* genes and orthogroups.

|  |  | Haplotype 1 | Haplotype 2 |
| --- | --- | --- | --- |
| Genes | Predicted gene candidates | 66,090 | 59,814 |
|  | **Genes in orthogroups** | 48,880 (74.0%) | 48,434 (81.0%) |
|  | **Number haplotype-specific genes** | 24,492 (37.1%) | 15,263 (25.5%) |
|  | **Number of shared genes** | 41,598 (62.9%) | 44,551 (74.5%) |
| **Orthogroups** | **Total number of orthogroups** | 39,959 |  |
|  | **Number of shared orthogroups** | 36,882 (95.4%) |  |
|  | **Haplotype-specific orthogroups** | 1,765 (4.6%) | 1,312 (3.4%) |
|  | **Number of orthogroups containing haplotype** | 38,647 (96.7%) | 38,194 (95.6%) |

### Gene functional annotation

To explore the potential functions of haplotype-specific and shared genes, all transcripts were functionally annotated. After choosing the best functionally annotated transcript for each gene based on the lowest e-value and highest score, we examined their COG (Clusters of Orthologous Groups) categories (Figure 3). Functional annotation successfully annotated 67.79% and 69.26% of genes for haplotype 1 and haplotype 2 respectively. Only 48.01% of all genes unplaced within an orthogroup were successfully functionally annotated, contributing the largest proportion to all non-functionally annotated genes. Genes not placed within an orthogroup and not functionally annotated may be false positives. Comparing COG categories of shared and non-shared genes revealed several categories containing different proportions of shared and haplotype-specific genes. These COG categories were found within genes associated with metabolism and information storage and processing, and also poorly characterised genes.

**Figure 3.**
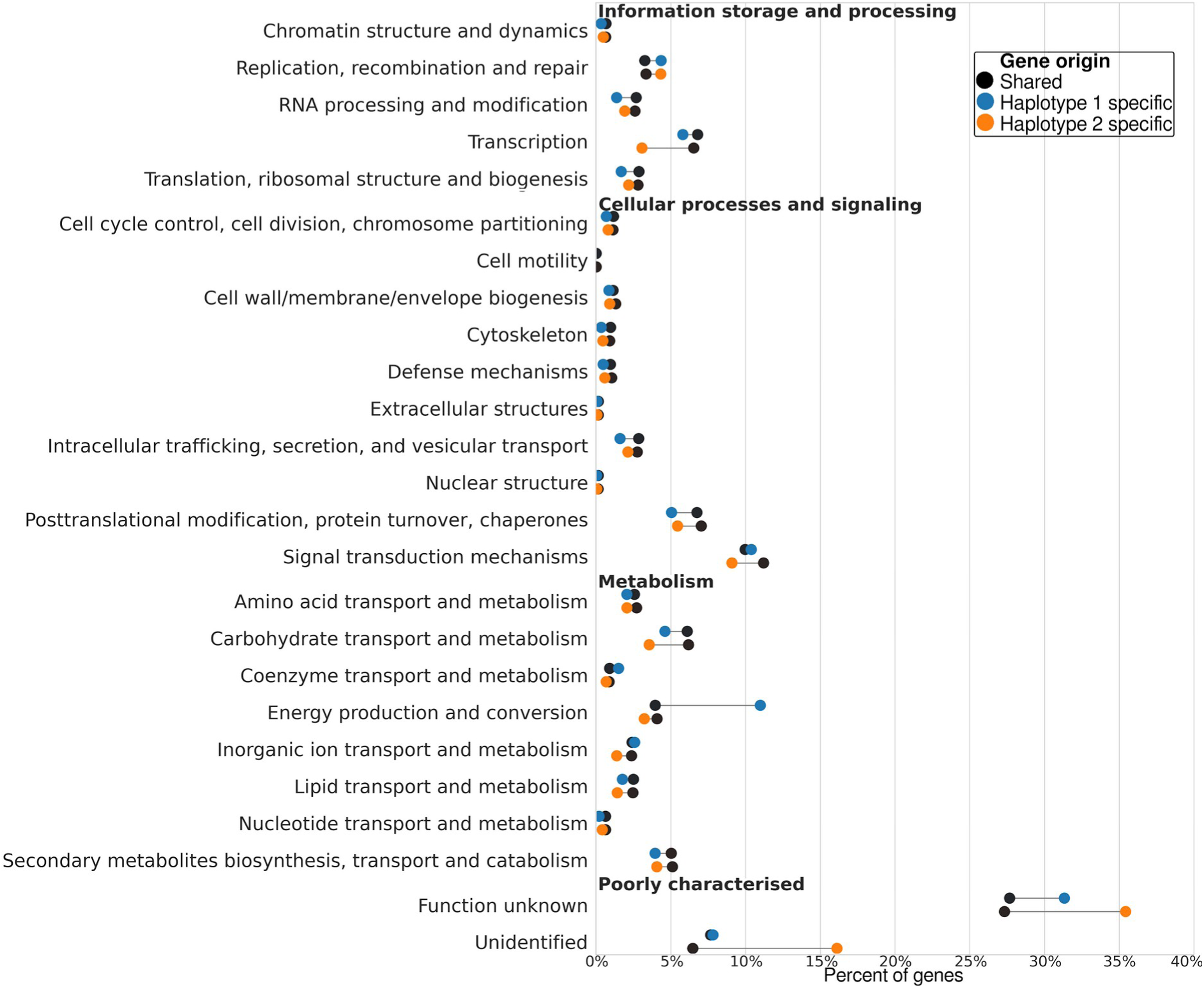
Investigating gene functions in both haplotypes. After functionally annotating all transcripts, the COG category of the best scoring transcript was chosen. All transcripts for both haplotypes were further categorised as shared, or haplotype-specific. Note: No genes were annotated as "General function prediction only".

### Genome synteny and structural variation between haplotypes

In addition to examining the gene differences between the two *E. regnans* haplotypes, we also examined the conservation of genome structure. After aligning the two haplotypes to each other, synteny, inversions, translocations, duplications, and haplotype-specific regions were annotated (Figure 4.A). On average, 77.7% of each haplotype was syntenic to the other haplotype, with the remaining proportion containing numerous structural variations. This included inversions (∼0.2% on average), translocations (∼8.8%), duplications (∼12.5%), or haplotype-specific regions (∼6.9%), representing insertions/deletions (Figure 4.D). Further examination of all genome regions revealed that syntenic regions are very large and very common, inversions are rare, translocations are moderately common, duplications are very common, and haplotype-specific regions are moderately common (Figure 4.C, 4.B).

**Figure 4.**
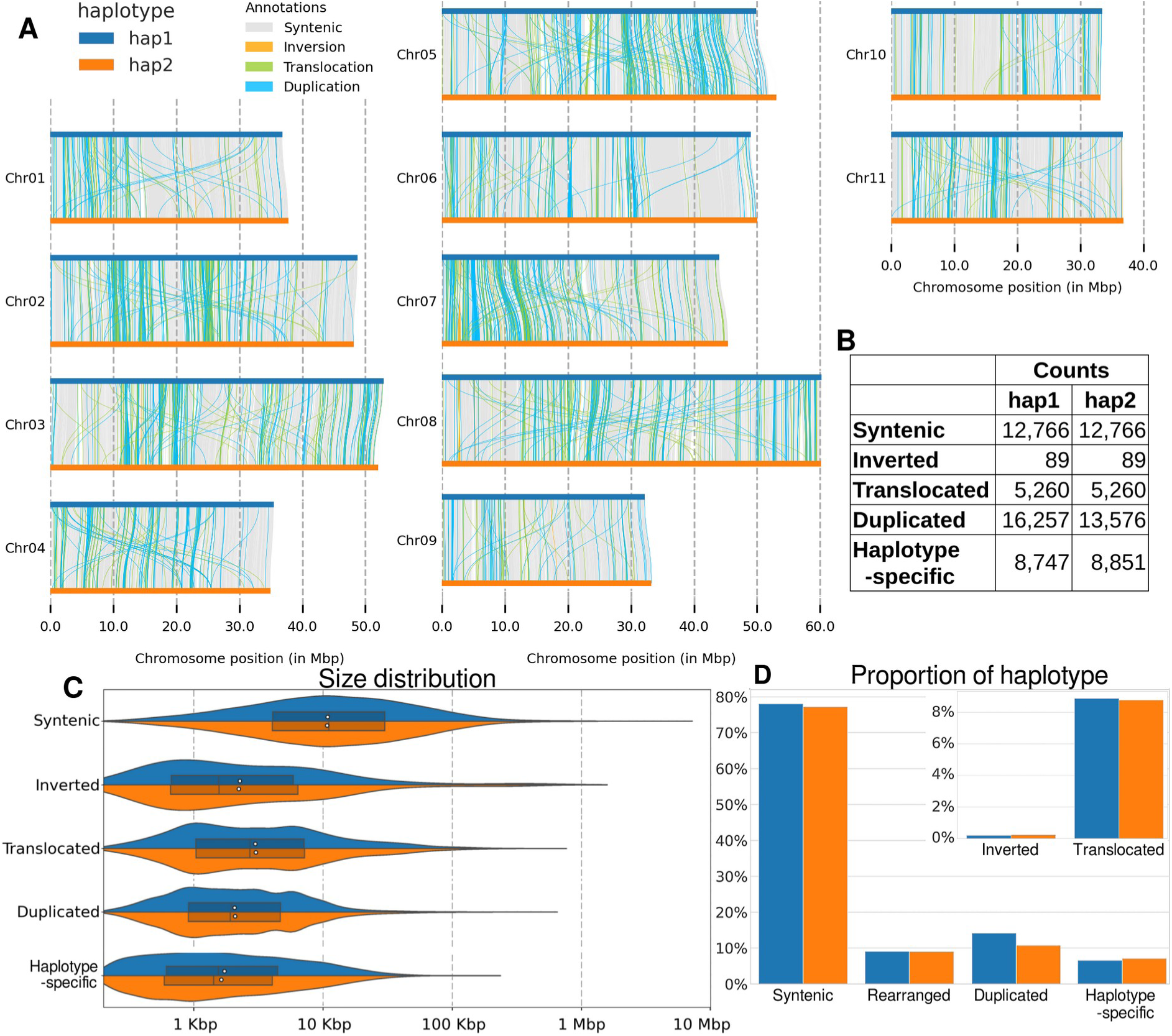
*E. regnans* genome synteny and structural variation. Haplotype 1 and 2 were aligned and all genome regions classified as syntenic, inverted, translocated, duplicated, or haplotype-specific. A) Karyotype plot shows locations of all genome regions, except inter-chromosomal translocations. B) The total number of each annotation type present within each haplotype. C) Size distribution of genome regions in both haplotypes. D) The proportion of each haplotype annotated as syntenic, haplotype-specific, inverted, translocated, and duplicated.

### Distribution of genes and TEs within across haplotypes

To examine the impact of structural variations on the genic content of each haplotype, the location of all genes was analysed. This analysis classified genes as originating in part of the genome that was syntenic, inverted, translocated, duplicated or haplotype specific. Similarly, the location of TEs were analysed, to determine if the inverted, translocated, duplicated or haplotype specific regions resulted from the movement, insertion, or deletion of TEs.

Analysis of genes revealed that the majority resided in syntenic regions, ∼87.34% of shared genes and ∼49.25% of haplotype-specific genes (Figure 5). Notably, the proportion of haplotype-specific genes within syntenic regions was significantly lower than that of shared genes. Shared genes outside of syntenic regions were predominantly found within duplications (∼8.21%), with a few found in translocations (∼4.23%). Inversions (∼0.11%) and haplotype-specific (∼0.11%) regions contained very few shared genes. Haplotype-specific genes were predominantly found within duplications (∼34.04%) and, to a lesser extent, translocations (∼16.34%), outside syntenic regions. This distribution significantly differed from shared genes, which were rarely found in these regions. Inversions (∼0.19%) and haplotype-specific (∼0.19%) regions again contained minimal shared genes. TE location analysis showed a similar trend, with the majority found within syntenic regions (∼70.62%). The remaining TEs were found predominantly in duplications (∼14.33%), haplotype-specific (∼7.98%) and translocated regions (∼6.86%). Inversions contained very few TEs.

**Figure 5.**
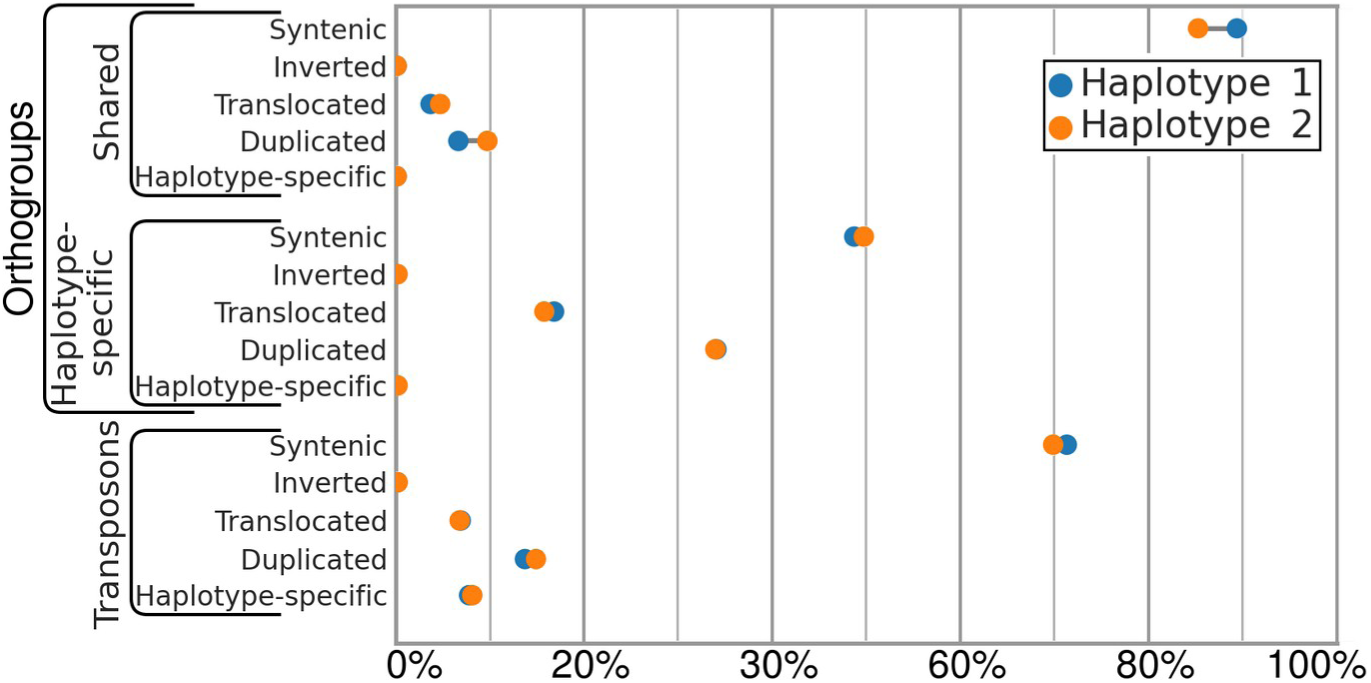
Comparison of the distribution of genes and TEs. All genes and TEs were classified based on their location within the genome (syntenic, inverted, haplotype-specific, duplicated, or translocated). Additionally, genes were categorised as shared (belonging to an orthogroup shared by both haplotypes) or haplotype-specific (belonging to a haplotype-specific orthogroup or unplaced).

## Discussion

### A chromosome-level, haplotype phased genome resource for *Eucalyptus regnans*

In this study, we generated high-coverage, long-read sequencing data, consisting of PacBio HiFi and ONT ultra-long, and chromosome conformation capture (Hi-C), to assemble a high-quality, haplotype-resolved genome for *E. regnans,* the Centurion. Achieving contig N50s > 36 Mbp per haplotype and 11 chromosome-scale scaffolds of N50s > 43 Mbp per haplotype, it is the most contiguous *Eucalyptus* genome to date, and only haplotype-phased genome [4, 23, 25]. The assembly was highly complete, achieving BUSCO completeness scores > 99% per haplotype. We identified 42 out of 44 expected telomeres, with each haplotype lacking only a single telomere, indicating a near T2T complete genome assembly. Long-read sequencing combined with advancements in *de novo* assembly algorithms, such as Hifiasm [17] and Verkko [26] has provided an unprecedented level of insight into genomes, by retaining haplotype sequence data that would typically be collapsed and/or removed in previous genome assembly pipelines. A critical advancement has been the recent integration of both long-read sequencing platforms (PacBio and ONT), and Hi-C data, in a single de novo assembler HiFiasm (UL) [17], utilised here in our study. This hybrid approach creates better genome assemblies by utilising both HiFi and ultra-long ONT reads to create a genome graph, and Hi-C data to improve haplotype phasing, resulting in highly contiguous, phased genomes.

### *Eucalyptus* genome architecture is shaped by structural variations

Investigation of the two haplotypes of *E. regnans* revealed ∼93.13% sequences were shared, and the remaining ∼6.87% was found to be haplotype-specific, likely originating from insertion and deletion polymorphisms. The shared sequences were highly syntenic (∼77.70%), but also had high levels of structural variations, including inversions (0.22%), translocations (8.84%), and duplications (∼12.48%). These structural variations were dramatically higher in abundance, size and distribution, in contrast to the highly syntenic haplotypes observed in agricultural crop genomes, such as rice [27], mango [28], grapes [29] and strawberry [30, 31]. Our findings align with a recent study across 33 *Eucalyptus* genomes that revealed an interplay of stable genome structure and accumulation of structural variations that drive genome divergence over time [4]. Similar results are being observed in other wild Myrtaceae family plants, such as Melaleuca [32]. However, the abundance of structural variations between the two haplotypes of a single *E. regnans* tree suggests a much greater degree of genetic variation within this species than previously assumed. This highlights the value of haplotype phased reference genomes and underscores the need for pan-genome approaches to capture the full spectrum of genetic diversity [33].

Comparative analysis of annotated genes revealed a high degree of conservation between haplotypes, with most of the orthogroups and genes found in both. However, a substantial number of genes were haplotype-specific. To investigate the origin of these unique genes, their locations were compared to syntenic, inverted, translocated, duplicated, and haplotype-specific regions. This analysis revealed a clear distinction: half of the haplotype-specific genes resided within structurally rearranged regions, primarily duplications, with a smaller proportion found in translocations. Conversely, shared genes were overwhelmingly located within syntenic regions. Similar to the analysis of genes, TE locations were examined to determine if structural variations between the haplotypes originated from TE movement. The majority of TEs resided in regions unaffected by inversions, translocations, duplications, insertions, or deletions. A small fraction was found within duplications, with an even smaller proportion residing in translocated and insertion/deletion regions. These findings suggest that while the core genome of *E. regnans* exhibits a high degree of synteny, extensive structural variations, particularly duplications, play a significant role in shaping the unique features of each haplotype. This dynamic genome structure, potentially fueled by TE induced recombination errors, may be a key factor underlying the remarkable adaptability of *Eucalyptus* to diverse environmental conditions [34, 35].

Placing all transcripts within COG categories indicated that the most significant difference in haplotype gene complements were in energy production and conservation, and transcription. This suggests that haplotype-specific variation contributes to environmental adaptation in *E. regnans*. Genes related to energy production and conservation could allow different individuals to thrive in different environmental niches [36, 37]. Similarly, variation in genes associated with transcription might enable them to respond to specific environmental signals by differentially regulating gene expression [38, 39]. These findings highlight the potential role of haplotype- or SV-specific genes in driving environmental adaptation

### An increasing need to conserve Australia’s giant *Eucalyptus* tree forests

Climate change is rapidly altering the environment worldwide, with increased intensity of drought, fire and floods [40–42]. This is having a significant impact on Australia’s iconic eucalypts, with extreme weather causing high tree mortality rates and dieback of forests [43]. Furthermore, unprecedented, mega-bushfires that occurred 2019-2020 caused widespread destruction of the natural landscape, especially eucalypt forests [8, 9]. This included *E. regnans* forests, and indeed the potentially record-breaking Centurion tree was partially burnt (Figure 1A), highlighting vulnerability of even the giants. Fires are particularly concerning for typically wet *E. regnans* forests, a keystone species with limited fire tolerance [44]. Unlike other eucalypts that can regrow vegetatively after fire (epicormic resprouters), *E. regnans* relies solely on seeds for regeneration (obligate seeder) [45], which take decades to become mature. Furthermore, tall trees such as *E. regnans* are huge stores of above and below-ground carbon (stem and root mass). Their loss would have catastrophic consequences for the forest ecosystem and loss of large carbon stores [46]. Loss of giant trees would greatly diminish the forest’s ability to annually sequester more atmospheric carbon, as these trees are still growing and gaining mass. The unique habitat provided by these forests would be substantially lost, impacting dependent wildlife populations and overall biodiversity [1]. For instance, tall, old growth *E. regnans* forests provide critical nesting sites and cavities (hollows) needed for a high biodiversity of birds and arboreal marsupials [47]. Therefore, protecting *E. regnans* forests and their towering giants becomes increasingly critical [48]. This study’s contribution lies in providing a high-quality genome of *E. regnans*, using samples from the Centurion itself. This resource will be instrumental in future research efforts aimed at understanding *E. regnans* population genomic diversity, complex growth traits including carbon capture and storage, and informing tall forest ecosystem conservation strategies.

## Conclusions

In this study, we assembled a high-quality, near T2T complete, haplotype-resolved diploid genome reference for *E. regnans*, the Centurion, a leading contender for the world’s tallest known flowering plant. This resource represents the most contiguous and complete reference genome for a *Eucalyptus* species, offering a foundation for future research into population genomics, functional genomics, and conservation of this ecologically significant tree species. Analysis revealed extensive structural variations and gene content differences between the two Centurion haplotypes, highlighting the remarkable genomic variation within *E. regnans*. Further exploration of this variation through pan-genomic or genome-graph approaches could provide deeper insights into the extent of the species’ genomic variation [33]. This is becoming tractable, given highly accurate long-reads from sequence consensus [49], specific base caller models [50], and deep learning error correction methods [51].

Sampling and sequencing additional trees across populations and performing genotype-environment associations can help uncover the molecular mechanisms of how *E. regnans* navigates variable environments, climates and potential threats like drought or fire [52]. This will lead to a better understanding of the genetic basis of environmental adaptation, carbon capture, and other key biological processes in *E. regnans*. Such knowledge is crucial for informing sustainable management practices and conservation efforts to protect carbon-dense giant tree forests and the unique ecosystems they support.

## Methods

### Sample collection and DNA extraction

The *Eucalyptus regnans* tree known as Centurion is located in the Huon Valley of Southern Tasmania, approximately 50 km SW of Hobart within the forestry estate (GPS -43.07708, 146.76859). Standing in an isolated small terrace of intact forest, surrounded by post-clearfell regeneration of *Eucalyptus*, it is thirty km south of the world’s tallest known flowering forest grove, the Tall Trees Reserve of the Styx Valley [53]. Tree climbing, by Giant Tree Expeditions, was required to sample leaf tissue. This leaf tissue was sent by local postal service with a cool pack to Australian National University, Canberra, where it was cryogenically stored in a -80°C until DNA extraction and Hi-C preparation.

High-molecular weight DNA was extracted with a magnetic bead-based protocol which is described in [54]. In brief, leaf material was ground with a mortar and pestle under liquid nitrogen, homogenate was washed with a sorbitol buffer, an SDS buffer lysis buffer was used followed by protein precipitation with potassium acetate. After binding the DNA to magnetic beads, it was washed multiple times with 80% ethanol before elution. DNA was size selected for fragments ≥ 20 kb using a BluPippin (Sage Science) for PacBio HiFi sequencing and ≥ 40 kb using a PippinHT (Sage Science) for ONT sequencing.

### Long-read sequencing of native DNA

For PacBio HiFi sequencing, the HMW DNA was sheared to approximately 18 kb fragments with a Megaruptor 3 (Diagenode), using 1 cycle 31x speed and 1 cycle at 32x speed. A PacBio SMRTbell library was prepared according to the manufacturer’s ‘SMRTbell Express Template Prep Kit 3.0’ (Pacific Biosciences). Sequencing was performed on a PacBio Revio 25M SMRT cell, with the circular consensus sequencing (CCS) mode to generate high-accuracy HiFi reads. DeepConsensus was automatically performed on the PacBio Revio, which increased sequencing accuracy [49].

For ONT sequencing, a native DNA sequencing library was constructed according to the manufacturer’s protocol ‘1D Genomic DNA by Ligation (SQK-LSK109)’. Sequencing was performed on an ONT MinION Mk1B devices using two FLO-MIN106D R9.4.1 flow cells. When sequencing declined (low active pore count, approximately 24 h), the flow cell was treated with DNAse I, primed again and more library was loaded, according to the manufacturer’s ‘Flow Cell Wash Kit (EXP-WSH004)’. This was performed at least twice to maximise total sequencing output of the flow cell, until the flow cell was expended. These reads were generated from as part of our previous study [4].

### Chromosome conformation capture with Hi-C

A proximity ligation library for chromosome conformation capture was created with a Phase Genomics Proximo Hi-C (Plant) Kit (version 4), according to the manufacturer’s instructions (document KT3040B). This kit utilised DpnII, HinFI, MseI, DdeI to digest the genome, sticky ends were then filled with biotin labelled nucleotides and the subsequent blunt ends were re-ligated to neighbouring molecules. The library was multiplexed with other projects and sequencing was performed on a NovaSeq 6000 (Illumina), using an S4 flow cell with a 300 cycle kit (150 bp paired-end sequencing).

### Assembly and scaffolding

All PacBio HiFi reads from the Revio were used in the assembly (≥ Q20). For ONT reads, both read ends were trimmed of 200 bp, followed by filtered to length of ≥ 1 kb and Q7, with NanoFilt (version: 2.8.0) [55]. For Hi-C reads, Illumina adapter sequences were removed (--nextera) and read pairs validated, using Trim Galore! (version: 0.6.10) [56].The *de novo* genome assembly was performed with Hifiasm ultra-long (UL) (version: 0.19.6-r595) [17], incorporating the PacBio HiFi, ONT ultra-long (--ul) and Hi-C reads (--h1 --h2). After assembly, both haplotypes were screened for contaminant contigs using BlobTools [57], which utilised minimap2 (version: 2.24) [58] and blast (version: 2.11.0) [59]. Subsequently both haplotypes were independently scaffolded using YaHS (version: 1.2a.2) [60] following the Arima Genomics mapping pipeline [61]. Briefly, bwa mem (0.7.17) [62] independently aligns both R1 and R2 Hi-C read sets to the current haplotype. Alignments were then filtered to remove chimeric reads and reads with poor MAPQ scores, subsequently the R1 and R2 alignment files were merged. Next, PCR duplicates were removed using Picard Tools (version: 2.26) [63]. Processed and combined pair-end alignments were then analysed with YaHS, generating a Hi-C contact map. As our ONT reads are very long and had high coverage, YaHS was run without the assembly error correction step. YaHS’s Hi-C contact map was checked, and when necessary, manually edited using Juicebox (version: 2.17) [64]. After manual curation, scaffolds were finalised with Juicer tools (version: 1.6) [65], producing chromosome-scale de novo genomes. Using tidk (version: 0.2.41) [66] candidate telomere sequences were generated and each candidate subsequently tested. BUSCO completeness analysis was calculated using compleasm (version: 0.2.2) [67].

### Annotation

De novo repeat libraries were generated for each haplotype using EDTA (version: 1.9.6) [68], including both simple repeats and transposable elements (TEs), and annotated with RepeatMasker (version: 4.0.9) [69]. The repeat-masked genomes were annotated for genes using BRAKER3 (version: 3.0.6) [70]. BRAKER3 aligned training proteins to our haplotypes using DIAMOND (version: 0.9.24) [71], and subsequently the ProtHint (version: 2.6.0) [72] pipeline generated the training data for AUGUSTUS (version: 3.5.0) [73]. Training protein sequences were obtained from the National Center for Biotechnology Information (NCBI) [24], including all available transcripts for Myrtaceae (Taxonomy ID: 3931) and Arabidopsis thaliana (Taxonomy ID: 3702).

Predicted gene candidates were subsequently organised into orthogroups using Orthofinder (version: 2.5.5) [74]. All candidate genes were functionally annotated for eggNOG orthogroup, COG category, GO term, KEGG term, and PFAM using eggNOG-mapper (version: 2.1.12; parameters: -m diamond --itype CDS --tax_scope Viridiplantae) [75].

### Genome alignments

We identified all shared sequences between our two *E. regnans* haplotypes through alignment using the MUMmer tool (version: 3.23) [76] with NUCmer (--maxmatch -l 40 -b 500 -c 200). NUCmer identified all shared 40-mers between genomes and merged adjacent 40-mers into a single alignment. Alignments were filtered to remove those < 200 bp and with an identity < 80% using MUMmer’s delta-filter tool. We chose a conservative 80% sequence identity threshold considering the high heterozygosity of *Eucalyptus* [3], and a higher score may incorrectly filter out real alignments. The filtered NUCmer alignments were then analysed for syntenic, inverted, translocated, and duplicated regions using SyRI (version: 1.6.3) [77]. All unaligned regions were annotated as haplotype-specific. A karyotype plot showing synteny and structural variations between haplotypes was created with Plotsr [78].

## Declarations

### Ethics approval and consent to participate

Not applicable.

### Consent for publication

Not applicable.

### Availability of data and materials

All raw sequencing data is available on the NCBI under the biosample accession number SAMN14929765. The genome generated for haplotype 1 is available at NCBI under bioproject accession number PRJNA1062543. Haplotype 2 is available at NCBI under bioproject accession number PRJNA1062542.

### Authors’ contributions

S.F. performed genome scaffolding, bioinformatic analysis and managed the data. A.J. performed DNA extractions, sequencing, *de novo* assembly and conceived the study. Y.D.B-N. performed fieldwork and sampling. All authors contributed to writing and review of the manuscript.

## Acknowledgements

We would like to thank the ACRF Biomolecular Resource Facility at the John Curtin School of Medical Research, ANU in Canberra, Australia, where PacBio Revio sequencing was conducted. This research acknowledges the support provided by NCRIS-enabled Bioplatforms Australia infrastructure.

Computational resources were provided by the Australian Government through the National Computational Infrastructure (NCI) under the ANU Merit Allocation Scheme. Additional resources were provided by the Australian National University Research School of Biology Dayhoff server.

Collection of *E. regnans* (the Centurion) was performed by Yoav D Bar-Ness as part of Giant Tree Expeditions. Collection and associated fieldwork was supported by the Tasmanian Museum and Art Gallery Jayne Wilson Bequest.

We also thank Dean Nicolle for supplying the species occurrence distribution map of *E. regnans* in Australia and his ongoing dedications to *Eucalyptus* trees.

We would like to acknowledge and thank Australia’s First Nations Peoples, the Traditional Custodians of the land on which *E. regnans* grows. We acknowledge and thank Elders past, present and emerging.

