## supplementary for "A diploid chromosome-level genome of *Eucalyptus regnans*: unveiling haplotype variance in structure and genes within one of the world’s tallest trees"

**Supplementary Table S1. Genome BUSCO scores.**

| | Haplotype 1 | | Haplotype 2 | |
|---|---|---|---|---|
| | **Count** | **Percent** | **Count** | **Percent** |
| Complete | 2,262 | 97.25% | 2,215 | 95.23% |
| Complete and single-copy | 1,938 | 83.32% | 1,924 | 82.72% |
| Complete and duplicated | 324 | 13.93% | 291 | 12.51% |
| Fragmented | 22 | 0.95% | 42 | 1.81% |
| Missing | 42 | 1.81% | 69 | 2.97% |

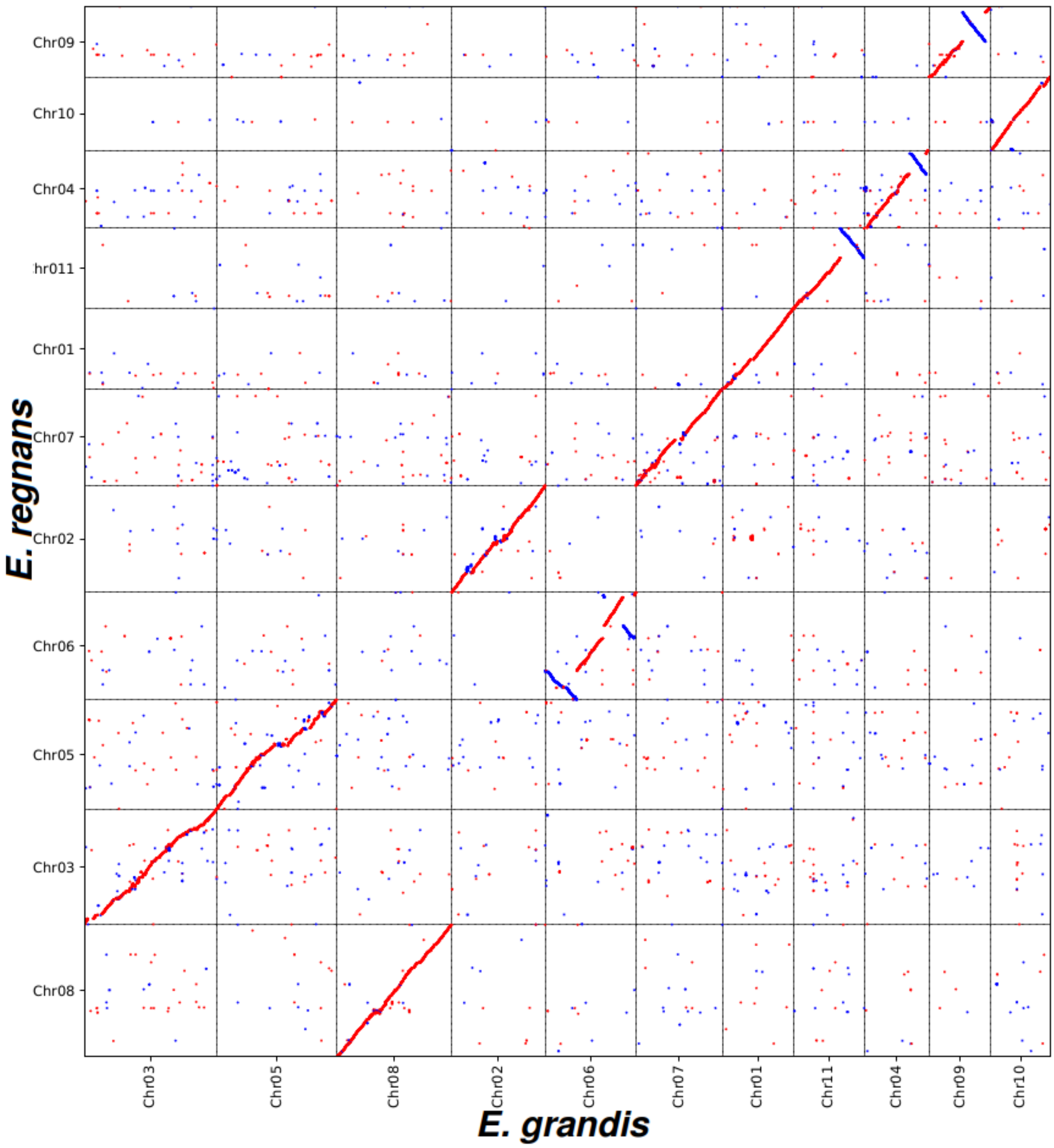


**Supplementary Figure S1.** ***E. regnans*** **haplotype 1 compared to** ***E. grandis***

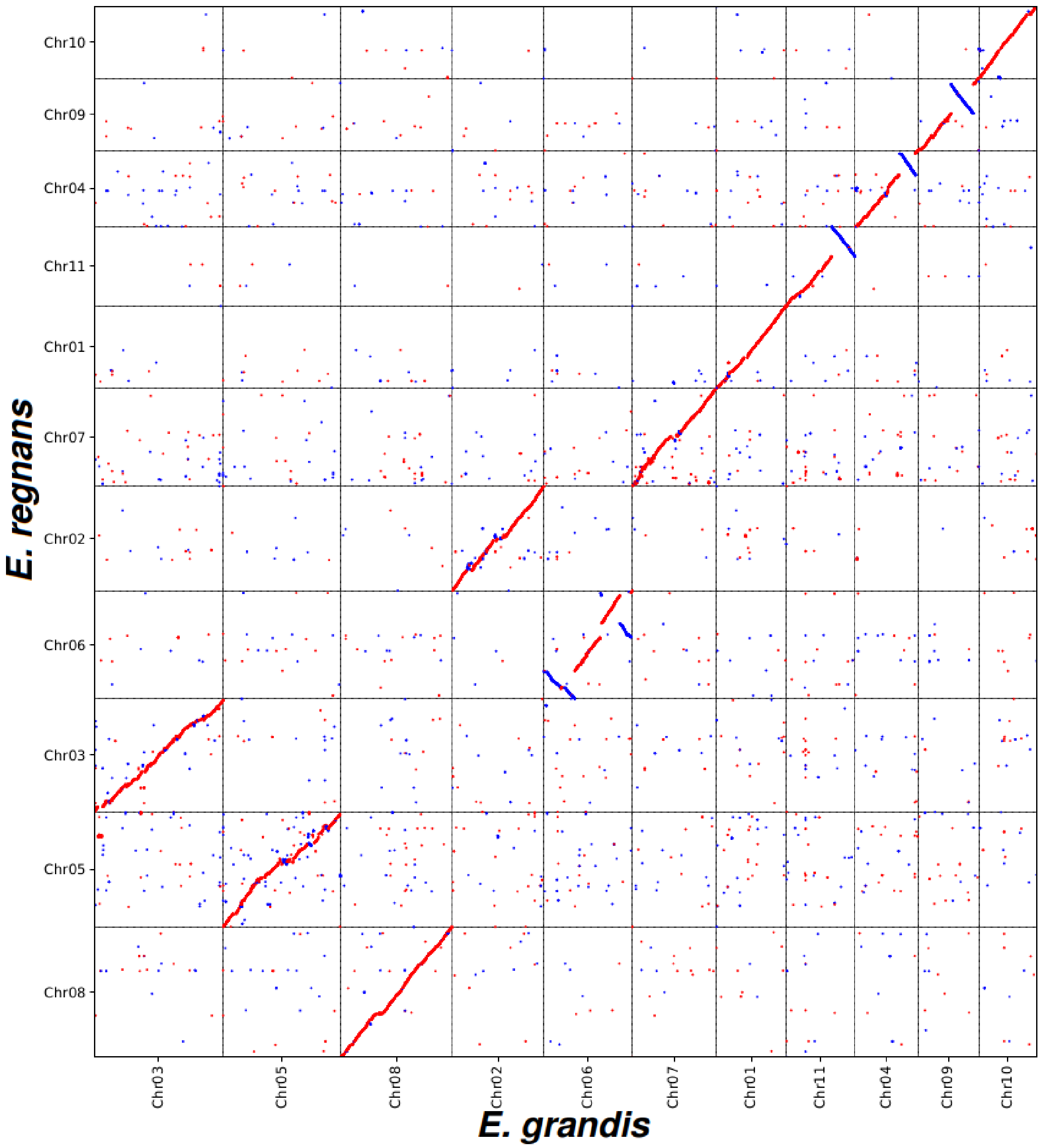


**Supplementary Figure S2.** ***E. regnans*** **haplotype 2 compared to** ***E. grandis***

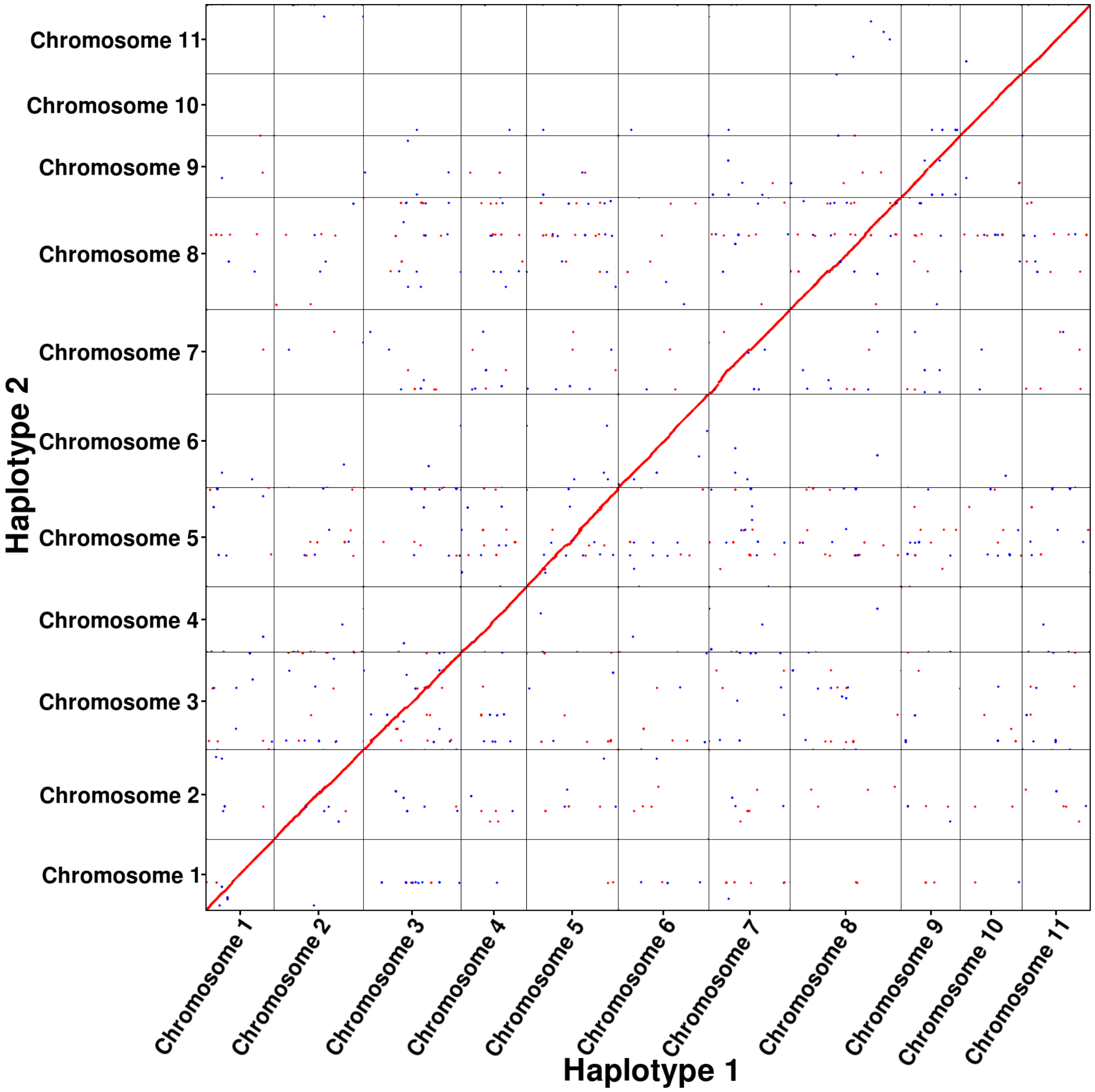


**Supplementary Figure S3.** ***E. regnans*** **haplotype 1 compared to haplotype 2.**
Confirmation that haplotypes were identically scaffolded.
